# Signatures of natural selection in the drug metabolizing enzyme genes: Opportunity for developing personalized and precision medicine

**DOI:** 10.1101/113514

**Authors:** Sheikh Nizamuddin, K. Thangaraj

## Abstract

Modern human experienced various selective pressures; including range of xenobiotics which contributed to heterogeneity of drug response. Many genes involve in pharmacokinetics and dynamics of drug, have been reported under natural selection. However, none of the studies have utilized comprehensive information of drug-centered PharmGKB pathways. We have extended this work and aimed to investigate sweep signals, using 1,798 subjects, from 53 Indian and 15 other world populations. We observed that modifiers which alters the biochemical function of other genes, have excess of natural selection (median std-z score=0.033±0.95; p-value=1.7×10^−9^-3.7×10^−3^). Taxane and statin primarily used for chemotherapy and lowering cholesterol level, respectively; and well known for heterogeneous drug response. We observed that pharmacokinetic pathway of taxane and statins are under natural selection (p-value=2.53×10^−9^and 2.73×10^−9^-1.09×10^−4^; q-value=1.28×10^−7^ and 6.91×10^−6^-1.1×10^−3^). We also observed signal of selection in Ibuprofen pharmacokinetics (p-value=1.76×10^−5^; q-value =2.22×10^−4^), beta-agonist/beta-blocker pharmacodynamics (p-value=4.79×10^−4^; q-value =4.04×10^−4^) and Zidovudin pharmacokinetics/dynamic pathway (p-value=7.0×10^−4^; q-value =5.06×10^−4^). Hard sweeps signals were observed in a total of 322 loci. Of which, 53 affect mRNA expression (p-value<0.001) and 16 were already reported with therapeutic response. Interestingly, we observed that Africans have experience 2 phases of natural selection, one at ~30,000 another at ~10,000 years before present.

## Background

Modern human migrated out of Africa ~75,000 years BP (before present) and colonized at diverse ecological conditions ^1-3^. During the migration, human populations have experienced various selection pressures, including range of xenobiotics, which were either present in new food sources or in new environment ^4^. Moreover, introduction of fire during the cultural revolution also had a major impact, as cooked food produce novel toxins and carcinogens ^5-9^. In order to cope up with different selection pressure, humans have acquired beneficial variations in the genome, including variation in the drug metabolizing enzyme gene(s) that are responsible for heterogeneous therapeutic response in the contemporary populations ^10^.

Many genes, which influence the drug level and their kinetics, have been designated as ADME (absorption, digestion, metabolism and excretion of drug) genes and categorized into core, extended, phase-I, phase-II and modifiers on the basis of their function; and were in natural selection among various populations ^10,11^. Several lines of evidences argue the existence of two type of selections; hard and soft sweeps ^12^. In hard sweep, advantageous mutation rise to higher frequency and single mutation event is sufficient to influence the phenotype; for example: *SLC24A5* mutation influences the skin pigmentation and *DARC* mutation influence the resistance against malaria^12-14^; while in soft sweeps, multiple advantageous mutations rise simultaneously and influence in polygenic manner; for example: cytokine-cytokine receptor signaling pathways ^15^. Although many studies on natural selection of ADME genes explored hard sweeps signals, soft sweeps signals remain unexplored. Hence, in the present study, we utilized the drug pharmacokinetics and dynamics pathway from PharmGKB database to explore the soft sweeps signals ^16^.

Several ADME genes that were reported in natural selection were found in out-of-Africa populations ^10^. But, it is not clear what happened to the populations those were left behind in Africa? To the best of our knowledge, none of the studies have revealed about the signals in pharmacogenomically important genes that were originated in Africa, after early human migration and colonization events. In the present study, we explored these ancient signals, using genome-wide markers.

## Results and discussions

### Evidence of natural selection

In this study, we explored overall signal of selection based on genome-wide genotype data (623,462 SNPs). Genic regions are considered more influenced under natural selection compared to non-genic regions and must be having high average std-z score. To explore it, initially we compared the whole genic SNPs with non-genic. Then, we compared genic SNPs of 873 pharmacogenomically important genes with non-genic, to understand that natural selection on pharmacogenomically important genes, are similar to whole genome genic region or not.

We observed significantly higher std-z score in genic region (299,317 SNPs; median = 0.0033) comparative to non-genic SNPs (324,145; median = −0.0033) (F = 1.0608, p-value <2.2×10^−16^; t = 9.8221, p-value < 2.2×10^−16^), which reflect that genic regions are under natural selection. To further explore that whether natural selection really acting on the genic region, variable range of flanking sequences (from 0 to 10^5^base pairs) around genes was considered (Table S3). SNPs that are far from genes are more towards non-genic designation and hence less std-z score is expected with increment of flanking region around gene. We observed that median std-z score of genic and non-genic SNPs decrease with increment of flanking region (**Figure 1C, 1D** and **Table S3**), except median score of non-genic increases at 2000 base pairs (bp). We also observed that per base-pair decrease in std-z score of genic region is less compared to non-genic regions, which is evident in **Figure 1E**.

**Figure 1.**
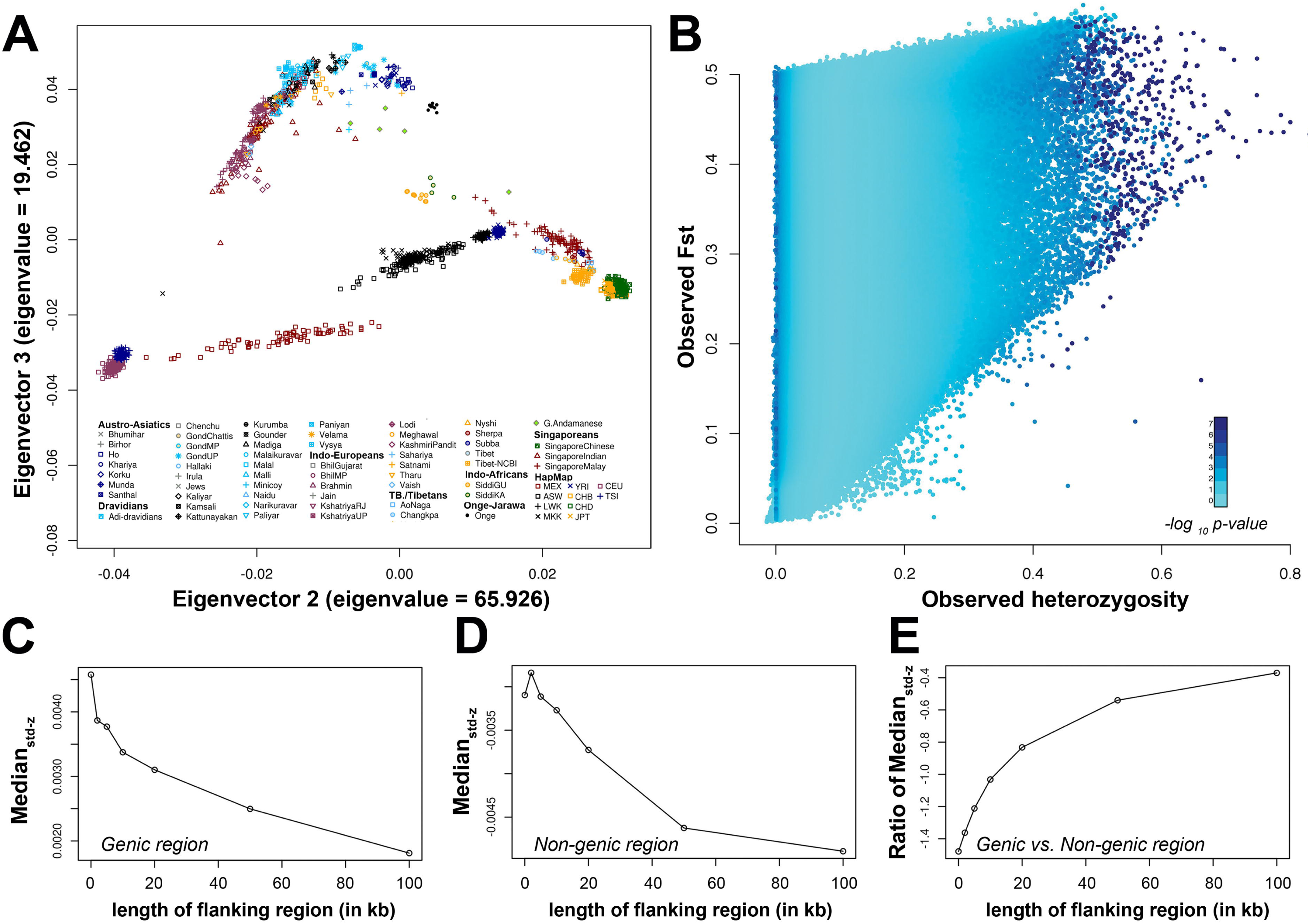
Simulation analysis using hierarchical island model, samples genetic clustering and signal of natural selection in whole genic and in pharmacogenomically important genes. **A.** Principal component analysis to explore genic cluster of samples prior to simulation analysis. **B.** Outcome of the simulation analysis with Hierarchical island model. Figure **C and D** represents decrease in the median std-z score with increment of flanking region while E represents the ratio of genic’s median std-z score to non-genic’s score.

Similar pattern was found in pharmacogenomically important genes. A total of 19,721 SNPs from 873 genes have significantly higher std-z score compared to non-genic SNPs (std-z score: F = 1.0406, p-value = 1.1×10^−4^; t = 3.455, p-value = 5.5×10^−4^), suggests that natural selection acted on these genes that are pharmacogenomically important.

### Modifiers ADME genes are most influenced under natural selection

To find out whether high frequency of natural selection among different categories of pharmacogenomically important genes, we have classified SNPs into core (494 SNPs), extended (3,900), phase-I (1,533), phase-II (767), modifier (285), transporter (1,805) and other pathway (15,333) genes; and compared them with other genic, non-genic and among themselves. Std-z score distribution of each category was compared for all combinations, using median and median absolute deviation (MAD) (**Table S4**). We observed that high frequency of modifier genes are under natural selection (std z-score = 0.033± 0.95) compared to other ADME categories, non-genic and other genic regions (**Table 1, S4 and S5**).

**Table 1.** Statistical significance of std-z score among different categories

| Category |  | F test |  | t test |  | Cat. | F test |  | t test |  | Cat. | F test |  | t test |  |
| --- | --- | --- | --- | --- | --- | --- | --- | --- | --- | --- | --- | --- | --- | --- | --- |
| A | B | F value | P value | t value | P value | A | F value | P value | t value | P value | A | F value | P value | t value | P value |
| All pathway genes | All pathway genes | - | - | - | - | Core | 0.871 | 0.03834 | -1.1268 | 0.2603 | PhaseII | 1.1749 | 0.00147 | 0.9105 | 0.3628 |
|  | All pharma genes | 0.9522 | 0.03904 | 1.006 | 0.3145 |  | 0.8294 | 0.006982 | -0.6871 | 0.4923 |  | 1.1188 | 0.03883 | 1.2746 | 0.2028 |
|  | Core | 1.148 | 0.03834 | 1.1268 | 0.2603 |  | - | - | - | - |  | 1.3489 | 0.0003036 | 1.4701 | 0.1418 |
|  | Extended | 0.9326 | 0.00498 | 0.77 | 0.4413 |  | 0.8124 | 0.002932 | -0.754 | 0.4511 |  | 1.0958 | 0.09586 | 1.1844 | 0.2365 |
|  | Genic | 0.9664 | 0.002129 | 0.4506 | 0.6523 |  | 0.8418 | 0.009184 | -1.0601 | 0.2896 |  | 1.1355 | 0.01067 | 1.0164 | 0.3098 |
|  | Modifier | 0.6441 | 1.91×10<sup>-08</sup> | -2.6178 | 0.009315 |  | 0.5611 | 2.16×10<sup>-08</sup> | -2.8536 | 0.004514 |  | 0.7568 | 0.003648 | -1.8904 | 0.05934 |
|  | Nongenic | 1.0251 | 0.02377 | 3.6562 | 0.0002567 |  | 0.8929 | 0.08499 | -0.466 | 0.6414 |  | 1.2045 | 0.0001641 | 1.6541 | 0.09852 |
|  | Other pathway genes | 0.995 | 0.7473 | -0.0994 | 0.9208 |  | 0.8667 | 0.03215 | -1.1502 | 0.2506 |  | 1.169 | 0.002142 | 0.8806 | 0.3788 |
|  | PhaseI | 0.9134 | 0.01452 | -0.1291 | 0.8973 |  | 0.7956 | 0.002313 | -1.0382 | 0.2995 |  | 1.0732 | 0.2545 | 0.6903 | 0.4901 |
|  | PhaseII | 0.8511 | 0.00147 | -0.9105 | 0.3628 |  | 0.7414 | 0.0003036 | -1.4701 | 0.1418 |  | - | - | - | - |
|  | Transporter | 1.1528 | 6.96×10<sup>-05</sup> | 3.9733 | 7.31×10<sup>-05</sup> |  | 1.0042 | 0.943 | 0.931 | 0.3521 |  | 1.3545 | 3.83×10<sup>-07</sup> | 2.8642 | 0.004249 |
| All pharma genes | All pathway genes | 1.0502 | 0.03904 | -1.006 | 0.3145 | Extended | 1.0723 | 0.00498 | -0.77 | 0.4413 | Transporter | 0.8674 | 6.96×10<sup>-05</sup> | -3.9733 | 7.31×10<sup>-05</sup> |
|  | All pharma genes | - | - | - | - |  | 1.021 | 0.504 | 0.1453 | 0.8845 |  | 0.826 | 1.92×10<sup>-06</sup> | -2.7896 | 0.005304 |
|  | Core | 1.2057 | 0.006982 | 0.6871 | 0.4923 |  | 1.231 | 0.002932 | 0.754 | 0.4511 |  | 0.9959 | 0.943 | -0.931 | 0.3521 |
|  | Extended | 0.9794 | 0.504 | -0.1453 | 0.8845 |  | - | - | - | - |  | 0.809 | 2.14×10<sup>-07</sup> | -2.843 | 0.004493 |
|  | Genic | 1.0149 | 0.4855 | -0.8866 | 0.3754 |  | 1.0362 | 0.1146 | -0.6305 | 0.5284 |  | 0.8383 | 3.04×10<sup>-07</sup> | -4.03 | 5.81×10<sup>-05</sup> |
|  | Modifier | 0.6765 | 1.53×10<sup>-06</sup> | -2.8056 | 0.00534 |  | 0.6907 | 6.27×10<sup>-06</sup> | -2.7533 | 0.006244 |  | 0.5588 | 4.28×10<sup>-12</sup> | -3.7201 | 0.0002334 |
|  | Nongenic | 1.0766 | 0.0004986 | 0.7171 | 0.4734 |  | 1.0992 | 2.41×10<sup>-05</sup> | 0.8672 | 0.3859 |  | 0.8892 | 0.0005856 | -2.8961 | 0.003824 |
|  | Other pathway genes | 1.0449 | 0.06782 | -1.0563 | 0.2909 |  | 1.0669 | 0.0101 | -0.8209 | 0.4117 |  | 0.8631 | 4.25×10<sup>-05</sup> | -3.9921 | 6.75×10<sup>-05</sup> |
|  | PhaseI | 0.9592 | 0.3167 | -0.6777 | 0.498 |  | 0.9794 | 0.6204 | -0.5606 | 0.5751 |  | 0.7923 | 2.05×10<sup>-06</sup> | -2.7709 | 0.005624 |
|  | PhaseII | 0.8938 | 0.03883 | -1.2746 | 0.2028 |  | 0.9126 | 0.09586 | -1.1844 | 0.2365 |  | 0.7383 | 3.83×10<sup>-07</sup> | -2.8642 | 0.004249 |
|  | Transporter | 1.2107 | 1.92×10<sup>-06</sup> | 2.7896 | 0.005304 |  | 1.2361 | 2.14×10<sup>-07</sup> | 2.843 | 0.004493 |  | - | - | - | - |
| Other pathway genes | All pathway genes | 1.0051 | 0.7473 | 0.0994 | 0.9208 | PhaseI | 1.0948 | 0.01452 | 0.1291 | 0.8973 | Modifier | **1.552** | **1.91×10<sup>-08</sup>** | **2.6178** | **0.009315** |
|  | All pharma genes | 0.957 | 0.06782 | 1.0563 | 0.2909 |  | 1.0425 | 0.3167 | 0.6777 | 0.498 |  | **1.478** | **1.53×10<sup>-06</sup>** | **2.8056** | **0.00534** |
|  | Core | 1.1539 | 0.03215 | 1.1502 | 0.2506 |  | 1.2569 | 0.002313 | 1.0382 | 0.2995 |  | **1.782** | **2.16×10<sup>-08</sup>** | **2.8536** | **0.004514** |
|  | Extended | 0.9373 | 0.0101 | 0.8209 | 0.4117 |  | 1.021 | 0.6204 | 0.5606 | 0.5751 |  | **1.448** | **6.27×10<sup>-06</sup>** | **2.7533** | **0.006244** |
|  | Genic | 0.9713 | 0.01343 | 0.5572 | 0.5774 |  | 1.058 | 0.1132 | 0.2655 | 0.7906 |  | **1.5** | **1.91×10<sup>-07</sup>** | **2.6785** | **0.007825** |
|  | Modifier | 0.6474 | 2.98×10<sup>-08</sup> | -2.6011 | 0.009769 |  | 0.7052 | 6.47×10<sup>-05</sup> | -2.4287 | 0.01564 |  | - | - | - | - |
|  | Nongenic | 1.0303 | 0.01018 | 3.5756 | 0.0003504 |  | 1.1223 | 0.001114 | 1.1983 | 0.231 |  | **1.591** | **1.70×10<sup>-09</sup>** | **3.017** | **0.002784** |
|  | Other pathway genes | - | - | - | - |  | 1.0893 | 0.02178 | 0.0889 | 0.9291 |  | **1.545** | **2.98×10<sup>-08</sup>** | **2.6011** | **0.009769** |
|  | PhaseI | 0.918 | 0.02178 | -0.0889 | 0.9291 |  | - | - | - | - |  | **1.418** | **6.47×10<sup>-05</sup>** | **2.4287** | **0.01564** |
|  | PhaseII | 0.8554 | 0.002142 | -0.8806 | 0.3788 |  | 0.9318 | 0.2545 | -0.6903 | 0.4901 |  | **1.321** | **0.003648** | **1.8904** | **0.05934** |
|  | Transporter | 1.1587 | 4.25×10<sup>-05</sup> | 3.9921 | 6.75×10<sup>-05</sup> |  | 1.2621 | 2.05×10<sup>-06</sup> | 2.7709 | 0.005624 |  | **1.79** | **4.28×10<sup>-12</sup>** | **3.7201** | **0.000233** |

We speculated that above observation could be biased by other SNPs, which were not in natural selection, but present within the genes. Hence, to prove the previous observation, SNPs that were in strong natural selection (std-z score >3) were only considered (reason for considering this cut-off value is explained in “Hard sweeps signals of natural selection”). Thus, 83 SNPs of ADME and 239 of other pathway genes were selected. It was observed that 27.27% modifier genes (6 out of 22) were under natural selection; while other categories ranges from 11.11% to 20.83% (**Table S5**). Among these, Phase-II genes (11.11%; 6 out of 54) were least influenced category. Moreover, the number of genes could also be biased by linkage disequilibrium (LD) among SNPs of different genes; hence, haplotype blocks of all populations were explored. Only rs3805322 (*ADH4*) and rs1042026 (*ADH1B*) were in the same haplotype block in HapMap-JPT, while they were in different blocks in other populations. Hence, it is clear that percentage of genes in each category is not the artifacts of LD (linkage disequilibrium).

It is interesting to note that modifiers influence the expression and alters the biochemical function of other ADME genes (www.pharmaadme.org). Alteration in the function of this category can cause major influence on pharmacogenomic system.

### Soft sweeps (polygenic adaptation) signals of natural selection

Much of selection acts as polygenic way (soft sweeps) with subtle influence on variation irrespective of hard sweeps, where either selection coefficient (>1%) or time of fixation is high^12^. To find the “soft sweep” signals, we utilized 102 pharmacogenomically important pathways (PharmGKB database).Of which, 45 pathways with less number of genes (< 10) were excluded from further analysis (**Table S6**). It is evident from previous reports that highest and median std-z score of SNPs assigned to gene have significant correlation and hence, highest score can be best representative of std-z score distribution within gene ^15^. Same is true in the present study, where a significant correlation (r = 0.436; p-value < 2.2×10^−16^; −0.448 95% C.I.) was observed between highest and median std-z score. Therefore, genic SNPs with highest std-z score were assigned as representative for the genes (total 16,594) in further soft-sweep signal analysis.

We observed multimodal distribution of std-z score, with 3 humps, having different range of score and density (**Figure 2A**). Maximum number of genes (11398: 68.69% of total gene) exists in the first hump (island-I) and have std-z score distribution from −2.378 to 0.824 (median = 0.54); intermediate number of genes (5031: 30.32 %) exist in second hump (island-II) and have distribution from 1.774 to 5.0 (median = 2.878); while in third hump (island-III), minimum number of genes (164: 0.99%) exists with std-z score from 5.1024 to 5.733 (median = 5.6091).

**Figure 2.**
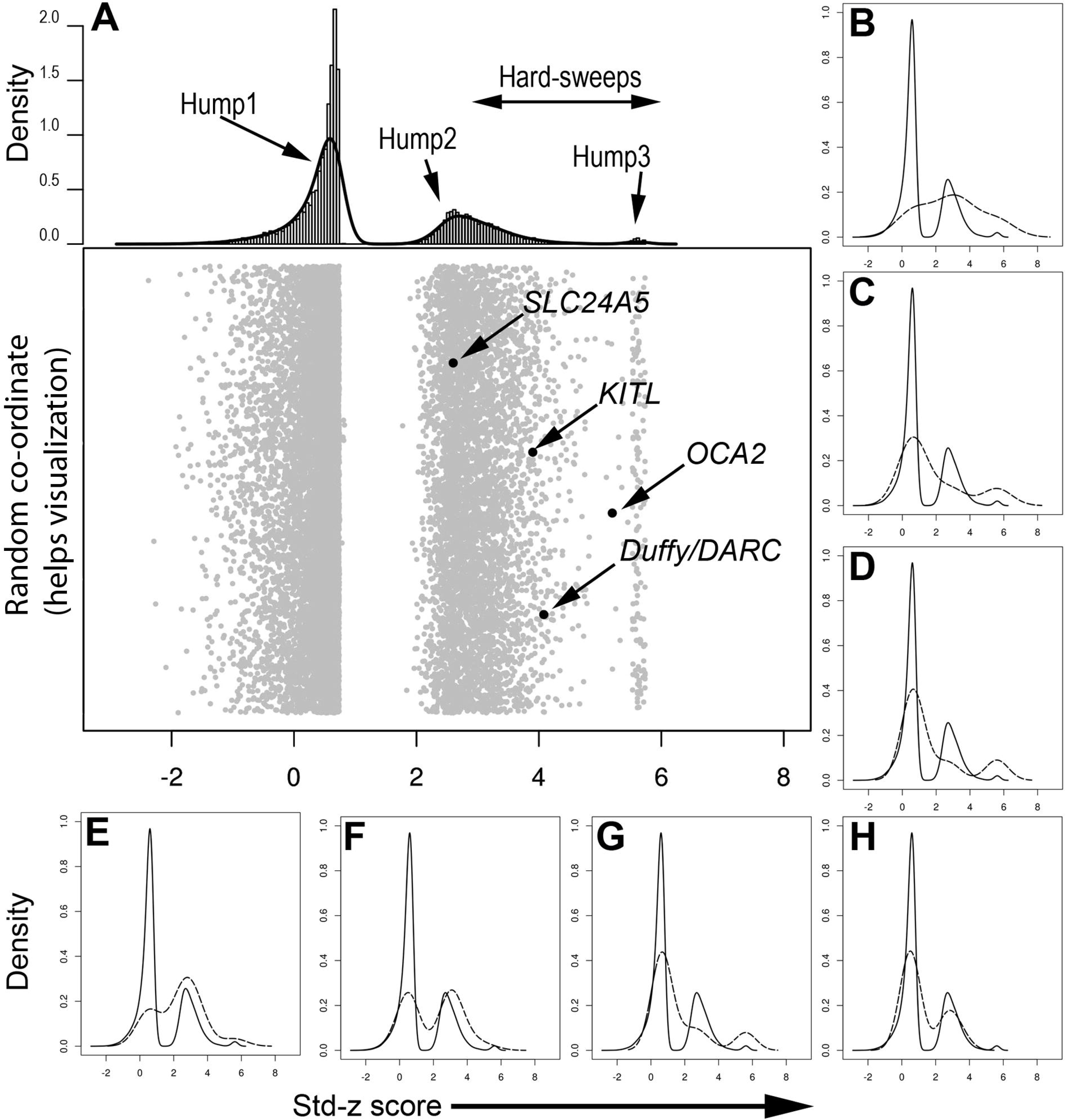
Soft sweep signals. In A, upper panel represents the density of std-z score where 3 humps can be observed while lower panel represents the distribution of std-z score of gene’s representative SNPs and few previously reported hard and soft sweep signals. The curve with dotted lines in **B, C, D, E, F, G** and **H** represent the distribution of std-z scores for Taxane pharmacokinetics, Fluvastatin pharmacokinetics, Atorvastatin-Lovastatin-Simvastatin pharmacokinetics, Statin generalized pharmacokinetics, ibuprofen pharmacokinetics, β-agonist β-blocker pharmacodynamics and Zidovudin pharmacokinetics/dynamics pathway’s genes respectively, while other curve with solid line in **B, C, D, E, F, G** and **H** represents the distribution of std-z scores for representative SNPs of all genes.

Further, we used the number of genes, present within 3 islands and compared them with genes of pathways, using simple chi-square test with 2 degree of freedom. For multiple test correction, p-values were corrected and converted into q-values (**Figure S3**) (false discovery rate control method).^17^ We have observed that 7 pathways were under natural selection (p-value < 7×10^−4^; q-value < 5.06×10^−3^), which includes pharmacokinetics pathway of Taxane, Fluvastatin, Atorvastatin-Lovastatin-Simvastatin, Ibuprofen and Statin; pharmacodynamic pathway of β-agonist/β-blocker; and pharmacodynamic and kinetics pathway of Zidovudin (**Table 2 and S7**) (**Figure 2B, 2C, 2D, 2E, 2F, 2G and 2H**). We speculated that these 7 pathways might be having the significant number of common genes which are under natural selection. But, we did not find the same (**Table S8**).

**Table 2.** Statistical significance of pathways. Three humps/islands were found in distribution curve of std-z score (Figure 3B). The corresponding number of the genes and percentage in each hump is given in the header of this table. Moreover, for each pathway, distribution of genes (number: percentage) in each hump is given with corresponding p-value and q-value.

| Pathway<sup>†</sup> | Total Genes | Hump1; (number of genes: percentage)<br>(11399: 68.69%) | Hump2; (number of genes: percentage)<br>(5031: 30.32%) | Hump3; (number of genes: percentage)<br>(164: 0.99%) | $\chi^2$p-value | q value |
| --- | --- | --- | --- | --- | --- | --- |
| TPP | 10 | 3: 30% | 5: 50% | 2: 20% | $2.53 \times 10^{-9}$ | $1.28 \times 10^{-7}$ |
| FPP | 12 | 8: 66.66% | 2: 16.67% | 2: 16.67% | $2.73 \times 10^{-7}$ | $6.91 \times 10^{-6}$ |
| A-L-SPP | 13 | 9: 69.24% | 2: 15.38% | 2: 15.38% | $8.79 \times 10^{-7}$ | $1.48 \times 10^{-5}$ |
| IPP | 16 | 11: 68.75% | 3: 18.75% | 2: 12.5% | $1.76 \times 10^{-5}$ | $2.23 \times 10^{-4}$ |
| SGPP | 19 | 10: 52.63% | 7: 36.84% | 2: 10.53% | $1.09 \times 10^{-4}$ | $1.10 \times 10^{-3}$ |
| BA/BBPP | 58 | 26: 44.83% | 31: 53.45% | 1: 1.72% | $4.79 \times 10^{-4}$ | $4.04 \times 10^{-3}$ |
| ZPDP | 17 | 5: 29.41% | 11: 64.71% | 1: 5.88% | $7.00 \times 10^{-4}$ | $5.06 \times 10^{-3}$ |
| <sup>†</sup>TPP: Taxane pharmacokinetics pathway; FPP: Fluvastatin pharmacokinetics pathway; A-L-SPP: Atorvastatin-Lovastatin-Simvastatin pharmacokinetics pathway; IPP: Ibuprofen pharmacokinetics pathway; SGPP: Statin generalized pharmacokinetics pathway; BA/BBPP: β-agonist/β-blocker pharmacodynamics pathway; ZPDP: Zidovudine pharmacokinetics and dynamics pathway | | | | | | |

Taxane and statin drugs are primarily used for chemotherapy and lowering cholesterol level. Taxane produced by the plants of the *Taxus* sp., while statins naturally produced by *Peniecillium* and *Aspergillus* fungi. Interestingly, statins are also observed in various natural sources including dairy products, whole grains, almonds and other nuts, fatty fish, pure sugar cane, apple cider vinegar, and many vegetables. On the basis of our results, we speculate that modern human experienced selection pressure in the past against taxane and statin like organic molecules, which are present in environment. Due to which beneficial mutations have increased only in those populations experienced in selective pressure; while in others frequency is less; and this can be the reason for different therapeutic responses in the present day populations. We presume that the same is true for Zidovudin and Ibuprofen.

### Hard sweeps signals of natural selection

The cut-off std-z score ≥ 3 was chosen to filter the SNPs in “hard sweeps” (Table S9). The best examples of “hard sweeps”: African *Duffy/DARC, SLC24A5, OCA2* and *KITLG,* ^12^ exists in hump2/Island-2 with std-z score 4.08, 2.6, 5.199 and 3.9 (**Figure 2A**). It justify our cut-off std-z score 3 for “hard sweep” signals. Frequency distribution of the 322 SNPs in the population, utilized in the present study, are given in **Figure S4**.

### Clinical annotation of hard sweeps signals through literatures

In the further step, we annotated the hard sweep signals with Ensembl-BioMart (version 75). Out of 322 SNPs, 16 (8 in ADME and 8 in other pathway genes) have been reported in association with drug-response (**Table S10**). Variant in the *ADH1B* (rs 1042026; missense variation: Arg47His) is well-known for its role in alcohol metabolism, had std-z score 3.14; ^18,19^ and previously reported under natural selection during Neolithic period ^20,21^. The rs363333 (*SLC18A2*: std-z score = 3.155) and rs892413 (*CHRNA6*; std-z score = 3.047), which are associated with tobacco and alcohol metabolism/dependence were also observed under selection ^22-25^.

The G allele of rs1056836 plays protective role against prostate cancer through metabolism of carcinogens including those ones, which are produced during meat cooking ^26-29^. We observed this SNP (rs1056836) under natural selection with std-z score 3.14. Moreover, we also observed high frequency of G allele in ‘out of Africa’ population (>0.558). East-Asians have very high frequency (0.92) of G allele and Europeans have less (0.558) among “out-of-African” population, while Africans have frequency ranging 0.124-0.357. Indo-Africans (Siddi) (0.3571) were close to African populations. Intriguingly, CC genotype is associated with decrease survival rate of prostate cancer patients, when treated with docetaxel compared to GC and GG genotypes ^30^. Natural selection is the reason for different frequency spectrum of these genotypes and hence, for heterogeneous drug response.

Another important variant, rs472660 (intronic) in *CYP3A43* is significantly associated with olanzapine clearance and having std-z score 4.32 ^31^. The AA genotype of rs472660 is well-known for the reason of racial differences in inefficacy and/or adverse reaction. Within Indian subcontinent, only Indo-African (Siddi) has AA genotype with 0.357 frequency, similar to other African populations (0.444-0.70%). GIH and Singapore-Indians (2.27 and 1.2, respectively) have similar frequency as European populations (CEU = 1.21 and TSI = 3.41). Interestingly, we found 2 SNPs, rs11103482 (*RXRA*; std-z score = 3.3) and rs2984915 (JUN; std-z score = 3.47),were under natural selection, which are known to influence measles vaccine immunity, an important finding which suggests that not only present day drug responses are outcome of natural selection, but also vaccine responses ^32,33^. *CYP3A4* is one of the important ADME enzymes, which metabolize ~50% of the drug and known to influence inter-individual responses. We found rs1851426 (std-z score = 5.61) in *CYP3A4* gene that influence phenotype variability with probe drug quinine ^34^. Similarly, rs2854450 (*EPHX1*) was found to be associated with diisocyanate-induced asthma was also in natural selection (std-z score = 3.07) ^35^.

Anti-psychotic drug Olanzapine is having higher rate of discontinuation, due to its inadequate response or hypersensitive reaction. The rs472660 (*CYP3A43*; AA genotype) has reported in association with clearance of Olanzapine, was observed under natural selection (std-z score = 4.32) ^31^. Frequency of AA genotype of rs472660 is higher in African population (44-70%), while Europeans (1.21-3.41%), HapMap-GIH (2.27%) and Singapore-Indians (1.2%) have significantly low frequency. Interestingly, the “AA” genotype was observed only in Indo-Africans (35.71%). We have also identified the variants that are associated with dose/response of anticoagulants (Phenoprocoumon: rs11150604: *STX4*: std-z score = 4.78), ^36^ resistant to anti-angiogenic therapies (Bevacizumab: rs9582036: *FLT1*: std-z score = 3.884) ^37-39^, response to psychological stress (Cortisol: rs242924: *CRHR1*: std-z score = 3.05) ^40-42^ and response in chemotherapy (Bortezomib and Vincristine: rs6457816: *PPARD*: std-z score = 4.0; Celecoxib: rs6017996: SRC: std-z score = 3.065; Sunitinib, Panitumumab, Oxaliplatin and Bevacizumab: rs9582036: *FLT1*: std-z score = 3.884) ^43-46^. Besides these, rs10059859 (*PDE4D*: std-z score = 5.59) was found to be associated with esophageal function and rs1403527 (*NR1I2*: std-z score = 3.36) was reported to be in worldwide differentiation, in “Hard sweeps” ^47,48^.

### Expression analysis

After clinically annotating the hard sweep signals, we explored the effect of these 322 SNPs on the mRNA expression. For this, we utilized whole genome expression data and genotype of HapMap individuals (details are given in method section); and observed 53 SNPs (19 in ADME and 34 in other pathway genes) affecting the mRNA expression with p-value < 0.001 (**Figure S5 and Table 3**). Of which, 3 SNPs (rs1056836, rs 11103482 and rs6457816) in ADME genes and 3 SNPs (rs2984915, rs9554316 and rs9582036) in other pathway genes have already been clinically annotated.

**Table 3:** Expression analysis of hard sweep signals

| Chr | rsID | Phys. Pos.<sup>+</sup> | An<sup>*</sup> | Dr<sup>*</sup> | M (A)<sup>‡</sup> | std-z score | Gene | Type | Probe-ID | p-value | Gene's pathways and drugs<sup>†</sup> | Reference |
| --- | --- | --- | --- | --- | --- | --- | --- | --- | --- | --- | --- | --- |
| 1 | rs2984915 | 59253695 | T | C | T | 3.4680616 | *JUN* | Other pathways genes | GI_44890066 | $6.04 \times 10^{-6}$ | VPP, EIPP, TCPP | 16 |
| 1 | rs6681761 | 165868887 | T | C | C | 3.2051442 | *UCK2* | Other pathways genes | GI_20357519 | $1.31 \times 10^{-4}$ | FPP | 16 |
| 2 | rs1056836 | 38298203 | C | G | C | 3.136306384 | *CYP1B1* | PhaseI: Extended | GI_13325059 | $4.78 \times 10^{-5}$ | APP; TPP | 16 |
| 2 | rs17023214 | 39225915 | C | G | G | 3.1178133 | *SOS1* | Other pathways genes | GI_15529995 | $6.75 \times 10^{-30}$ | VPP; EIPP | 16 |
| 2 | rs7571608 | 39325132 | G | A | G | 3.0668693 | *SOS1* | Other pathways genes | GI_15529995 | $1.05 \times 10^{-25}$ | VPP; EIPP | 16 |
| 2 | rs7564481 | 39330377 | G | A | A | 3.1706149 | *SOS1* | Other pathways genes | GI_15529995 | $3.98 \times 10^{-28}$ | VPP; EIPP | 16 |
| 2 | rs3791358 | 101031865 | A | G | A | 3.078068634 | *CHST10* | PhaseII: Extended | GI_20127466 | $5.17 \times 10^{-15}$ | - | - |
| 3 | rs12632779 | 196009099 | T | A | A | 3.5206544 | *PCYT1A* | Other pathways genes | GI_31543384 | $8.06 \times 10^{-4}$ | LPP | 16 |
| 4 | rs6446647 | 4445315 | A | G | A | 3.6628182 | *STX18* | Other pathways genes | GI_39725935 | $3.59 \times 10^{-4}$ | NPP | 16 |
| 4 | rs11097642 | 99929768 | T | C | C | 3.103200082 | *METAP1* | PhaseI: Extended | GI_24308008 | $4.03 \times 10^{-5}$ | - | - |
| 4 | rs1230178 | 99935882 | G | A | A | 3.704828454 | *METAP1* | PhaseI: Extended | GI_24308008 | $2.83 \times 10^{-4}$ | - | - |
| 4 | rs7683532 | 99944057 | A | G | G | 3.107445521 | *METAP1* | PhaseI: Extended | GI_24308008 | $6.6 \times 10^{-5}$ | - | - |
| 4 | rs4698804 | 110940045 | T | C | C | 3.5641027 | *EGF* | Other pathways genes | GI_8393298 | $1.39 \times 10^{-7}$ | VPP; EIPP | 16 |
| 4 | rs4698804 | 110940045 | T | C | C | 3.5641027 | *EGF* | Other pathways genes | GI_42659252 | $2.63 \times 10^{-5}$ | VPP; EIPP | 16 |
| 4 | rs4698804 | 110940045 | T | C | C | 3.5641027 | *EGF* | Other pathways genes | GI_20149595 | $3.1 \times 10^{-4}$ | VPP; EIPP | 16 |
| 4 | rs4698804 | 110940045 | T | C | C | 3.5641027 | *EGF* | Other pathways genes | GI_22095396 | $4.21 \times 10^{-4}$ | VPP; EIPP | 16 |
| 4 | rs4834424 | 115553989 | G | A | G | 3.388034316 | *UGT8* | PhaseII: Extended | GI_40254470 | $7.30 \times 10^{-4}$ | - | - |
| 4 | rs6831964 | 115589159 | T | C | C | 3.188523115 | *UGT8* | PhaseII: Extended | GI_40254470 | $7.35 \times 10^{-4}$ | - | - |
| 4 | rs6815181 | 115595647 | C | T | T | 3.207004849 | *UGT8* | PhaseII: Extended | GI_40254470 | $7.35 \times 10^{-4}$ | - | - |
| 4 | rs4834427 | 115600000 | A | G | A | 3.180948098 | *UGT8* | PhaseII: Extended | GI_40254470 | $3.45 \times 10^{-4}$ | - | - |
| 5 | rs6870785 | 125905569 | A | G | G | 3.33501115 | *ALDH7A1* | PhaseI: Extended | GI_4557342 | $2.85 \times 10^{-21}$ | CT | 60 |
| 5 | rs7736031 | 125905992 | T | C | C | 3.244859299 | *ALDH7A1* | PhaseI: Extended | GI_4557342 | $2.02 \times 10^{-20}$ | CT | 60 |
| 6 | rs9658100 | 35356640 | G | T | G | 4.604327323 | *PPARD* | Modifier: Extended | GI_29171748 | $4.07 \times 10^{-6}$ | TDR | 61 |
| 6 | rs6457816 | 35362848 | C | T | C | 4.002995311 | *PPARD* | Modifier: Extended | GI_29171748 | $1.05 \times 10^{-5}$ | TDR | 61 |
| 6 | rs6906237 | 35375526 | A | C | A | 3.17958232 | *PPARD* | Modifier: Extended | GI_29171748 | $9.28 \times 10^{-5}$ | TDR | 61 |
| 7 | rs1029951 | 151245740 | C | T | C | 3.17667474 | *PRKAG2* | Other pathways genes | GI_33186924 | $6.5 \times 10^{-4}$ | MPP | 16 |
| 8 | rs1383888 | 31758989 | T | C | C | 3.10383996 | *NRG1* | Other pathways genes | GI_7669521 | $4.31 \times 10^{-4}$ | EIPP | 16 |
| 8 | rs7821196 | 32084242 | C | T | T | 4.02468353 | *NRG1* | Other pathways genes | GI_4758525 | $4.14 \times 10^{-4}$ | EIPP | 16 |
| 8 | rs7821196 | 32084242 | C | T | T | 4.02468353 | *NRG1* | Other pathways genes | GI_7669521 | $4.42 \times 10^{-4}$ | EIPP | 16 |
| 9 | rs11103482 | 137287860 | C | T | C | 3.300724399 | *RXRA* | Modifier: Extended | GI_21536318 | $2.36 \times 10^{-5}$ | DD | 62 |
| 10 | rs3737180 | 31607361 | C | T | C | 3.75270214 | *ZEB1* | Other pathways genes | GI_28077090 | $1.53 \times 10^{-11}$ | DPP | 16 |
| 10 | rs7894459 | 31611256 | T | C | T | 3.83044223 | *ZEB1* | Other pathways genes | GI_28077090 | $5.09 \times 10^{-13}$ | DPP | 16 |
| 10 | rs161279 | 31684697 | C | G | C | 3.5887484 | *ZEB1* | Other pathways genes | GI_28077090 | $1.74 \times 10^{-11}$ | DPP | 16 |
| 10 | rs161258 | 31710631 | A | G | G | 3.10648723 | *ZEB1* | Other pathways genes | GI_28077090 | $1.40 \times 10^{-9}$ | DPP | 16 |
| 10 | rs161272 | 31745639 | A | G | A | 3.7803372 | *ZEB1* | Other pathways genes | GI_28077090 | $4.33 \times 10^{-12}$ | DPP | 16 |
| 10 | rs161275 | 31772766 | T | C | T | 3.48646289 | *ZEB1* | Other pathways genes | GI_28077090 | $2.29 \times 10^{-11}$ | DPP | 16 |
| 10 | rs1761379 | 31783338 | T | C | T | 3.39061409 | *ZEB1* | Other pathways genes | GI_28077090 | $2.05 \times 10^{-11}$ | DPP | 16 |
| 10 | rs220073 | 31808203 | C | A | C | 3.54653944 | *ZEB1* | Other pathways genes | GI_28077090 | $2.29 \times 10^{-11}$ | DPP | 16 |
| 10 | rs7906892 | 106025990 | T | G | T | 3.073267976 | *GSTO1* | PhaseII: Extended | GI_4758483 | $2.56 \times 10^{-7}$ | PBC | 63 |
| 13 | rs9554316 | 28881335 | T | G | T | 4.0877333 | *FLT1* | Other pathways genes | GI_32306519 | $3.97 \times 10^{-9}$ | SPP; VPP | 16 |
| 13 | rs9582036 | 28885408 | C | A | C | 3.8837962 | *FLT1* | Other pathways genes | GI_32306519 | $9.41 \times 10^{-11}$ | SPP; VPP | 16 |
| 13 | rs7981680 | 28893870 | G | C | G | 4.0565268 | *FLT1* | Other pathways genes | GI_32306519 | $1.32 \times 10^{-11}$ | SPP; VPP | 16 |
| 13 | rs2093821 | 28969209 | C | T | C | 3.048008 | *FLT1* | Other pathways genes | GI_32306519 | $9.48 \times 10^{-9}$ | SPP; VPP | 16 |
| 14 | rs9805889 | 24417399 | C | G | G | 3.452804966 | *DHRS4* | PhaseI: Extended | GI_32483356 | $1.09 \times 10^{-19}$ | XCRCG | 64 |
| 14 | rs8010151 | 24423582 | G | A | G | 3.410793528 | *DHRS4* | PhaseI: Extended | GI_32483356 | $1.73 \times 10^{-18}$ | XCRCG | 64 |
| 14 | rs10137990 | 24425058 | G | T | T | 3.523457569 | *DHRS4* | PhaseI: Extended | GI_32483356 | $5.42 \times 10^{-20}$ | XCRCG | 64 |
| 14 | rs11158346 | 61928306 | T | C | T | 3.0025389 | *PRKCH* | Other pathways genes | GI_28557780 | $1.30 \times 10^{-6}$ | AD | 16 |
| 16 | rs6500567 | 4083105 | G | A | G | 3.0060239 | *ADCY9* | Other pathways genes | GI_4557258 | $2.40 \times 10^{-5}$ | PPIP, BA/BBPP | 16 |
| 16 | rs12103309 | 4100268 | G | T | G | 3.0758123 | *ADCY9* | Other pathways genes | GI_4557258 | $1.32 \times 10^{-5}$ | PPIP, BA/BBPP | 16 |
| 17 | rs2159141 | 11981961 | C | G | G | 3.1247165 | *MAP2K4* | Other pathways genes | GI_24497520 | $1.70 \times 10^{-5}$ | TCPP | 16 |
| 17 | rs12944877 | 63035254 | G | A | G | 3.6272293 | *GNA13* | Other pathways genes | GI_31343475 | $1.25 \times 10^{-8}$ | PAIPP | 16 |
| 17 | rs1533075 | 63039373 | A | G | G | 3.5154491 | *GNA13* | Other pathways genes | GI_31343475 | $1.38 \times 10^{-6}$ | PAIPP | 16 |
| 17 | rs11656396 | 63046076 | C | T | T | 3.052676 | *GNA13* | Other pathways genes | GI_31343475 | $1.30 \times 10^{-8}$ | PAIPP | 16 |
| 17 | rs4791242 | 63057651 | T | C | T | 3.6692325 | *GNA13* | Other pathways genes | GI_31343475 | $5.82 \times 10^{-9}$ | PAIPP | 16 |
| 17 | rs1076094 | 73344711 | T | C | T | 3.0621962 | *GRB2* | Other pathways genes | GI_24431995 | $3.3 \times 10^{-4}$ | VPP; EIPP | 16 |
| 17 | rs4789188 | 73397871 | C | A | A | 4.1817875 | *GRB2* | Other pathways genes | GI_24431995 | $4.13 \times 10^{-4}$ | VPP; EIPP | 16 |
| 20 | rs819147 | 32889704 | C | T | C | 3.0766003 | *AHCY* | Other pathways genes | GI_9951914 | $4.41 \times 10^{-12}$ | TTP | 16 |

+Physical position are based on hg19 reference panel
\*An: Ancestral allele; Dr: Derived allele; Alleles matching with Chimpanzee are considered as ancestral while other as derived
‡Minor allele: In FigureS5 genotype of minor allele is represented as AA while major allele's genotype is represented as BB
†VP: Vemurafenib pharmacodynamics; EIP: EGFR inhibitor pharmacodynamics; TCP: Tacrolimus/cyclosporin pharmacodynamics; FP: Fluoropyrimidine pharmacokinetic; AP: Amodiaquine; TP: Taxane pharmacokinetics; LP: Lamivudin pharmacokinetics/dynamics; NP: Nicotine pharmacodynamics; CT: Cisplatin toxicity; TDR: Thalidomide and Docetaxel response; MP: Metformin pharmacodynamics; DD: Docetaxel deposition; DP: Diuretics pharmacodynamic; PBC: Platinum based chemotherapy-survival rates; SP: Sorafenib pharmacodynamics; XCRCG: Xenobiotics containing reactive carbonyl group; AD: Antidepressants; PPIP: Proton-pump inhibitor; BA/BBPP: β-agonist/ β-blocker pharmacodynamics; PAIP: Platelet aggregation inhibitor pharmacodynamics; TP: Thiopurine pharmacodynamic/kinetics

### Signal of ancient positive natural selection

Late Pleistocene and Holocene age of modern human experienced high ecological and demographic changes; and rise of agriculture. To find the acceleration of selective sweeps, we calculated age of variants (only for ADME genes) based on the length of extended haplotype homozygosity (EHH) on either side of derived allele, in East-Asians (JPT and CHB), Europeans (CEU), Mexicans (MEX), African (YRI), Indians (Indo-Europeans and Dravidians), Tibeto-Burmans & Tibetans, Onge and Siddi. We observed that in all populations, variants have split into 2 cluster: in the 1^st^ cluster SNPs have high range of age and std-z score (< 0.824) while in 2^nd^ cluster SNPs having less range with std-z score (> 1.774) (**Figure S6**). Since, our aim was to find the acceleration of natural selection, we proceeded further with the 2^nd^ cluster (std-z score>2). Further, we considered only those variants having age of < 60ky and estimated kernel density with 100 breakpoints. In all 8 populations analyzed, we found high density of SNPs near to ~ 10,000 year BP (before present).

Interestingly, we observed African population has 2 phase of acceleration (Figure 3). First phase of acceleration in this population was ~30,000 years BP while second phase was ~10,000 (same as other population). We did not find signal for natural selection at ~30,000 YBP in other populations suggesting that acceleration of natural selection in Africans might have happened after out-of-Africa migration. To understand this event, we explored the signals in Onge and Siddi populations. Onge is a “Negrito” population, who are the descendent of the first “out-of-Africa” population and inhabited in isolation for tens of thousands years in the Andaman and Nicobar islands of Indian sub-continent ^49^ while Siddi population is the recent (~400 years) migrant from Africa and has admixed with Indian populations ^50^. If natural selection would have acted after “out-of-Africa” migration, Onge populations must be carrying recent signals; but not the ancient signals due to their isolation while others African and Siddi population should be having both. Presence of signals only in Siddi not in Onge suggests that African continent possess 2 wave of acceleration compared to populations from rest of the world. Further, we tried to explore the difference between these 2 phases and found that the differences exist only in SNP level, not genes level. This is a preliminary observation and needs to be explored further.

**Figure 3.**
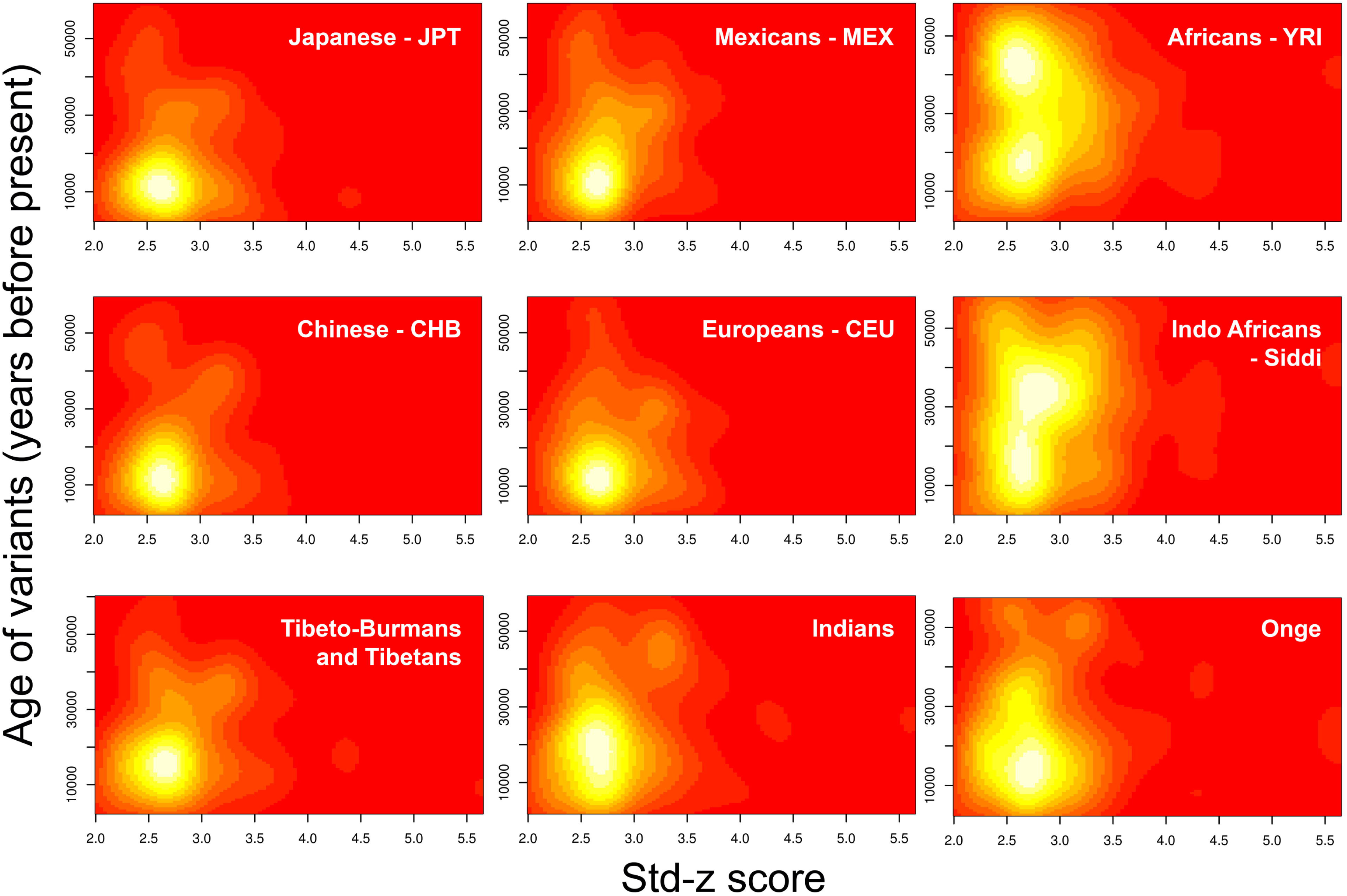
Evidence of two phase of natural selection in Africans and its comparison to other world populations.

## Conclusions

We identified signature of natural selection in pharmacogenomically important genes. Of which, modifiers which alters the biochemical function of other genes, have excess of natural selection. Using comprehensive information of drug-centered pathway from PharmGKB, we observed 7 pharmacokinetic and dynamic pathways under natural selection. Our study also identified 322 hard sweep signals. We have demonstrated that 53 affect mRNA expression level. In future, these SNPs can be utilized to understand heterogeneous drug response, personalized therapy in contemporary populations.

## Methods

### SNP data

A total of 53 Indian populations, who belongs to diverse social, linguistic and geographical background (4 Tibeto-Burman, 1 Tibetan-refugees, 12 Indo-Europeans, 22 Dravidians, 10 Austro-Asiatic, 2 Indo-Africans, 1 Great-Andamanese and 1 Onge) were selected for this study. The source of genotype data for these 53 Indian populations, are given in **Table S1**. In addition, 11 populations from HapMap, 3 from Singapore genome diversity project and 1 Tibetan population from NCBI-GEO database were also included (**Table S1**). This study was approved by the Institutional ethical committee of CSIR-Centre for Cellular and Molecular Biology, Hyderabad, India.

Further, 811 genes from 102 pharmacokinetic and pharmacodynamic important pathways have been selected from PharmGKB database ^16^. Of which, 31 were core ADME genes (8 transporter, 12 phase-I and 11 phase-II) and 92 were extended ADME genes (27 transporter, 39 phase-I, 13 phase-II and 13 modifier). Besides these, 1 core and 165 extended ADME (40 transporter, 72 phase-I, 42 phase-II and 11 modifier) genes from PharmaADME database, which are not in pathway, were also been selected. SNPs, which are within 10,000 base pairs (bps) of above described 977 genes were considered as genic and extracted from whole genome data after imputation. In quality control filtering (imputation R2<0.95), genotype information of 104 genes (including 5 ADME genes on X-chromosome) were excluded (**Figure S1**) and analyzed using the remaining 20,225 SNPs.

### Imputation of missing genotypes

We have already demonstrated that imputation of missing genotypes is more accurate with Indian haplotype reference panel ^51^. Hence, we utilized 56 Indian-specific haplotype (http://www.ccmb.res.in/bic/database_pagelink.php?page=snpdata) as reference panel for Indo-European, Dravidian and Austro-Asiatic; and performed imputation with Beagle-v3.3.1. Of which, only 623,462 SNPs having R2> 0.9, were selected for further analysis. For Tibeto-Burman and Tibetan, we utilized phased haplotype reference panel of JPT, CHB and CHD HapMap populations while for Siddi population, we utilized haplotype reference panel of YRI.

### Principal component analysis (PCA)

To genetically cluster the populations for coalescence simulations with hierarchical island model (discussed in subheading “Test for natural selection”), we performed PCA with EIGENSOFT package ^52^. Since, populations can differentiate on spurious axis due to linkage disequilibrium (LD) between SNPs, we excluded 193,014 SNPs with r^2^>0.75 prior to PCA and utilized remaining 444,610 SNPs. Further, we explored the clusters on initial 3 eigenvectors, having highest eigenvalues.

### Test for natural selection

To detect the signal of selection, population differentiation measured by F_*st*_, were used, because it gives collective information about all populations in single statistics in comparison to others *i.e.* HaploPS, XP-CLR, XP-EHH, his ^13,15,53-55^. However, absolute value of F_*st*_ could be misleading because of its correlation with heterozygosity ^56^, hence, p-value for F_*st*_ was assessed. For this, we generated empirical distribution of F_*st*_ with respect to their heterozygosity in 100,000 coalescence simulations with hierarchical island model (10 groups and 100 demes per group) ^57^ As proposed earlier that in this model, demes within the same group (continent) are assumed to exchange migrants at a higher rate than demes in different groups, which reflect the hierarchical nature of human continental regions and appropriate to use in simulation. To perform this simulation, we clustered the samples in 10 groups on the basis of their linguistic affiliations and genetic clustering observed in principal component analysis (PCA) (**figure 1A**); 1) Onge, 2) Great-Andamanese, 3) Austro-Asiatic, 4) Dravidians, 5) Indo-Africans, 6) Indo-Europeans, 7) Tibeto-Burman and Tibetans, 8) HapMap-Africans, 9) Singapore-Malay, Singapore-Chinese and HapMap-East Asians; and 10) Admixed populations with Indian ancestry(GIH and Singapore-Indians), HapMap-Europeans and HapMap-Mexicans.

Moreover, to understand the differentiation not only between populations but also between pharmacogenomically important genes and other genic (within 10,000 base pair of gene) & non-genic regions, above distribution was used to calculate the p-value for all 623,462 autosomal SNPs. It is evident in **Figure 1B** that SNPs having both low and high F_*st*_ value are under natural selection. Since, the aim of the study is to find pharmacogenomically important SNPs which are highly differentiated among populations and are under natural selection, we converted p-value into z-score, in a way that highly differentiated SNPs (having high F_*st*_ value) score high positive value ^15,57^ The qnorm function of R with Epanechnikov kernel density was used, for conversion of p-value to z score.

We observed that 623,462 SNPs are not equally distributed in genome (**Figure S2**). In this scenario, genomic regions with high density of SNPs are more likely to possess extreme value of z-score, only by chance and hence, standardization is needed. We split the whole genome into 13,933 fragments (each 2×10^5^ base pairs) and clustered in 20 bins according to their SNP density (**Table S2**). Further, z-score distribution within bin was used for standardization as proposed by Daub, *et al.* (2013) ^15^. It is briefly, explained below:

Suppose, an *x* locus has z score “*z_x_*” and on basis of SNP density belongs to *i^th^* bin, then standardized z-score (std z-score) for *x* will be;

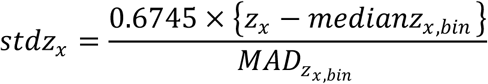

Where; median absolute deviation,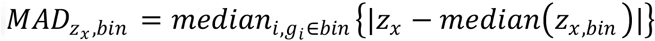

To represent the central tendency of the distribution, median has always been used in place of mean; while for representing the statistical dispersion, median absolute deviation (MAD) divided by 0.6745 was used in place of standard error/deviation, as these values are more realistic even in case of long tail or exponential distribution. Interestingly, if distribution is symmetrical, median becomes equal to mean. Hence, if, we do not know the pattern of distribution, median is the best choice. All statistical analysis *i.e.* F test, t test, pathway significant analysis was performed with R basic package.

### Clinical annotation of Hard-sweep signals

In order to explore the clinical significant of naturally selected loci, Ensembl-BioMart (Ensembl version 75: February 2014) was used to extract literatures in which hard-sweep loci obtained in the present study, was associated with the drug responses.

### Expression analysis

To find the significance of hard sweep signals, we also utilized whole genome expression dataset of HapMap subjects from Gene Expression Omnibus (GEO) with GSE6536 ID and genotype data of same individuals from ftp://ftp.ncbi.nlm.nih.gov/hapmap/genotypes/2009-01_phaseIII/plink_format. For the expression of each mRNA, we used whole genome normalized expression score and performed Wald-test for quantitative trait association analysis with Plink software ^58^.

### Age of variants

To calculate the rough estimate of age of variants, we used the formula mention by Zhernakova, A. *et. al* ^59^. For this, initially, we calculated extended haplotype homozygosity score *x* on both side of core allele with “rehh” package of R. The alleles, which are matching with the Chimpanzee are considered as ancestral while other as derived allele. Only the age of derived alleles were calculated and considered as core allele. Suppose the genetic distance between left and right side of core SNP is *r,* the generation time *G* can be calculated as:

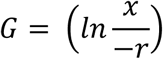

(Block)

For the calculation of age, we consider only those genetic distance where x is equal to 0.25. In the present study, we consider that 1 generation is equal to 25 years.

## Declarations

### Ethics approval and consent to participate

Collection of DNA samples and use of genetic data, were approved by ethical committee of CSIR-Centre for Cellular and Molecular Biology, India.

### Consent for publication

Not applicable.

### Availability of data and material

The datasets generated during and/or analyzed during the current study are not publicly available due to data and privacy protection considerations but may be available on justified request.

## Competing interests

Authors declare that they have no competing interests.

## Funding

This work was supported by CSIR Network project—EpiHeD (BSC0118), Government of India. Sheikh Nizamuddin was supported by ICMR JRF-SRF research fellowship.

## Authors’ contributions

KT was project leaders. SN analyzed data. SN and KT wrote the manuscript. All authors read and approved the final manuscript.

## Acknowledgements

Sheikh Nizamuddin acknowledges ICMR for JRF-SRF research fellowship.

